# Natural Selection in a Synthetic Yeast Endosymbiont Promotes Stable Coexistence

**DOI:** 10.1101/2024.07.31.606091

**Authors:** Lubica Supekova, Han Zhou, Izabella H. Barcellos, Catherine Nguyen, David A. Dik, Peter G. Schultz

## Abstract

Bacteria engulfment by a higher order host is believed to be the beginning of an evolutionary process that ultimately formed mitochondria. In an effort to experimentally elucidate the early effects of natural selection on bacteria resident in a eukaryotic host, a synthetic endosymbiont model system has been exploited. Here we describe a reproducible series of mutations that were observed after *Escherichia coli* was passaged within *Saccharomyces cerevisiae* for >8 passages which led to enhanced coexistence of bacteria within the yeast. These naturally selected mutations, formed by gene acquisition of transposable elements in *rcsC, cpxA*, and *idnK*, result in both functional and non-functional protein products with phenotypic effects on the bacteria that promote endosymbiont stability.

## INTRODUCTION

The endosymbiotic theory^1^ describes a series of events that began with engulfment of bacteria by an archaeon host cell, followed by extensive genetic alterations, and a corresponding phenotypic transformation that culminated in the modern mitochondria.^2^ Because these events occurred over a billion years ago, our understanding of this process is largely limited to phenotypic and genotypic analyses of mitochondria from phylogenetic trees. The bacterial origin of mitochondria has not formally been identified, but several bacteria have been proposed including Gram-negative purple non-sulfur bacteria,^3^ alphaproteobacteria Rickettsiales,^4^ and more recently, Gram-negative marine Iodidimonas.^5^ The evolutionary prerequisites for the formation of a mitochondrion from a bacterium include significant genomic reduction, loss of the cell wall, re-targeting of proteins to and from the bacterium, and horizontal gene transfer between the bacterium and host to promote coexistence.^6, 7^ The ultimate validation of this proposed series of events would be recapitulation of the evolutionary steps in a synthetic model system.^8^

In a previous report, we designed and implemented a model system to study endosymbiotic evolution in the laboratory using the bacterium *Escherichia coli* (*E. coli*) and eukaryotic yeast *Saccharomyces cerevisiae (S. cerviseae)*.^9^ *E. coli* is an ideal model bacterium as it shares genetic features conserved in Gram-negative bacteria and is highly amenable to recombinant manipulation both genetically and environmentally. *E. coli* deficient in NAD^+^ due to gene ablation of *nadA*, were engineered to export adenosine triphosphate (ATP) via an adenosine diphosphate (ADP)/ATP translocase^10^ to provide energy to the yeast host. In parallel, mitochondrial defective yeast *S. cerevisiae* were prepared by disruption of the cytochrome c oxidase subunit 2 (*COX2*) gene (strain NB97), rendering the strain incapable of growth in media with a non-fermentable carbon source.^11^ We found that it was also necessary to express SNARE-like proteins in *E. coli* to prevent cytosolic clearance of the bacteria.^12^ A polyethylene glycol (PEG) fusion protocol was adapted from a mitochondrial transformation technique^13^ to permit fusion of *E. coli* Δ*nadA* into *S. cerevisiae* NB97, forming endosymbionts capable of surviving up to 4 passages (>40 doublings), after which the yeast host cells released bacterial endosymbionts. This loss of codependence with passaging suggests that other as of yet unidentified factors affect endosymbiont stability i.e., stable coexistence. Nonetheless, these synthetic endosymbionts permitted a preliminary study of bacterial genome reduction within a yeast host.^14^ Herein, we report a set of natural mutations acquired by the bacterium during residence in yeast that promote enhanced endosymbiont stability.

## RESULTS AND DISCUSSION

### Discovery of Evolved Bacterium 1

To understand those genetic factors that affect bacteria and host codependence, *E. coli*-yeast endosymbionts were generated and extensively passaged on agar plates to identify any mutations that might promote coexistence. Endosymbionts that survived eight passages (1 passage every three days, >80 doublings) were inoculated in liquid endosymbiont growth media containing 1 M sorbitol, 3% glycerol, 0.1% glucose and 50 µg/mL carbenicillin. Endosymbionts are not stable under these liquid conditions and release bacteria. The bacteria were collected and analyzed by whole genome sequencing. Transposon insertion sequences^15^ were uniformly discovered in three genes, *cpxA*::IS1 family transposon, *idnK*::IS4-like element, and *rcsC*::IS4-like element. This strain is hereafter referred to as Evolved Bacterium 1 (EB1). Wild-type *E. coli* survives in this liquid media confirming that any mutational effects are not simply caused by the endosymbiont growth media. In addition, these mutations are not observed when *E. coli* are serial passaged in sorbitol.^16^

### EB1 Exhibits Enhanced Endosymbiont Stability

We next evaluated if EB1 forms a more stable endosymbiont, defined by *E. coli* retention in yeast over time, by widefield imaging. Endosymbionts of EB1 and ΔnadA were prepared, and the percent of yeast hosting intracellular bacteria was determined by fluorescent microscopy after four consecutive passages, each spanning three days. The number of fluorescent yeast cells, indicating the presence of intracellular bacteria, were counted at passage one and four.

Consistent with the observation that EB1 forms more colonies after fusion, by microscopy 25.9% of yeast fused with EB1 were GFP positive, while only 3.9% of yeast fused with ΔnadA retained intracellular bacteria (Figure 1A). After four passages 9.6% of yeast fused with EB1 maintain intracellular bacteria, compared to only 1.1% of yeast fused with the ΔnadA strain (representative images are provided in Figure 1B and 1C). Note, endosymbionts harboring EB1 displayed a uniform reduction in size. Collectively, the quantitative fluorescence microscopy findings indicate that the EB1 genotype leads to a significant increase in the number of endosymbionts upon fusion and after extensive rounds of replication.

**Figure 1.**
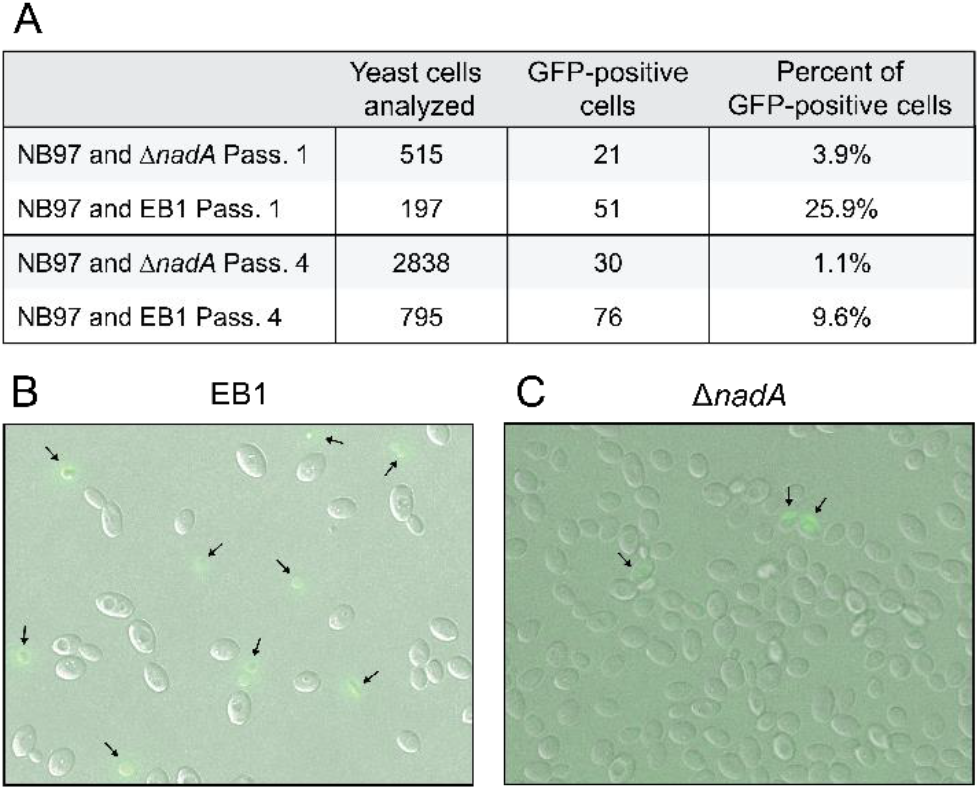
(A) Metrics of endosymbiont image analysis to determine the percent of GFP-positive yeast (NB97) cells hosting either the Δ*nadA* or EB1 endosymbionts in passage 1 and 4. Both *E. coli* strains genomically encode *gfp*. Representative overlaid brightfield and fluorescent images of (B) EB1 and (C) Δ*nadA* endosymbionts at passage 4. Arrows mark confirmed fluorescent yeast cells.

### Transposon Acquisition in *rcsC, cpxA* and *idnK*

A new preparation of yeast-*E. coli* endosymbionts was prepared and the state of each gene after each passage was monitored by colony PCR to assess a potential increase in gene size over time due to transposon insertion, and to confirm that the three transposons were acquired while *E. coli* inhabited yeast. Indeed, in surviving endosymbionts transposon insertions into *cpxA, idnK*, and *rcsC* began to appear at passage three, and by passage five >95% of observed PCR products indicated a transposon insertion for each gene (Figure 2A). The discovery of EB1, encoding three gene disruptions, was reproducible in replicate experiments. Surprisingly, DNA sequencing of PCR products revealed that these transposon insertions all occurred at the same sites in independent experiments suggesting that they may modulate rather than abolish the activity of the encoded proteins.

**Figure 2.**
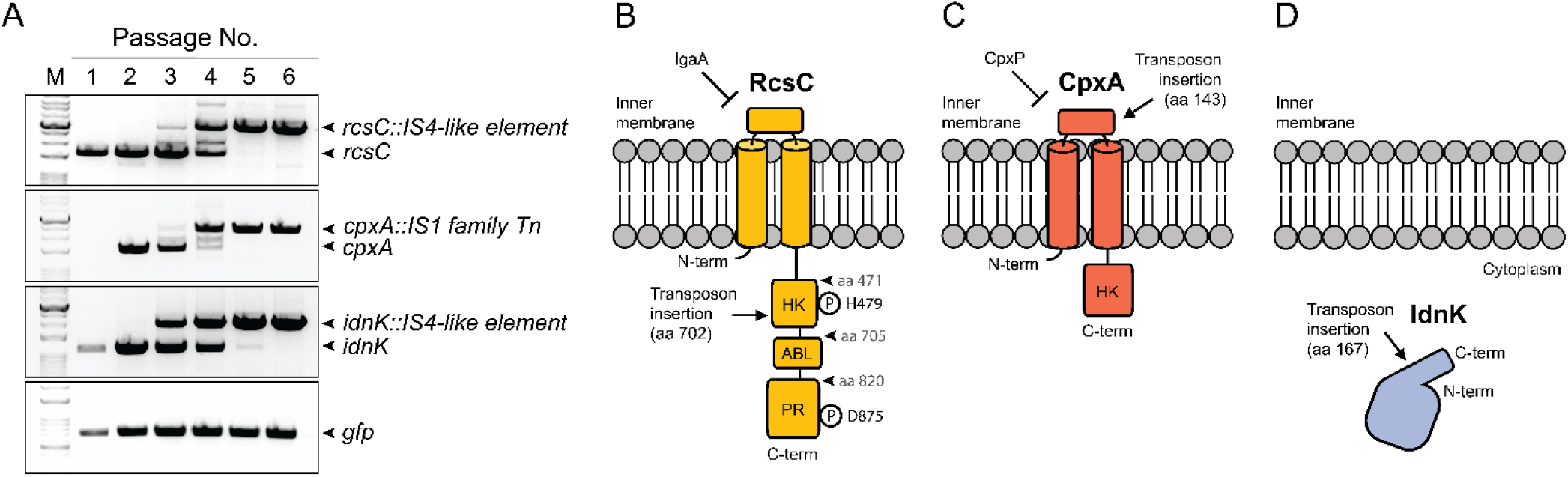
Identification and validation of the EB1 genotype. (A) Agarose gel of a PCR product for analyzing the acquisition of transposon insertions by *rcsC, cpxA*, and *idnK* is shown for each passage. PCR of the green fluorescent protein gene (*gfp*) on the bacterial genome is provided as a loading control. Depiction of (B) RcsC, (C) CpxA, and (D) IdnK protein domain architectures and transposon insertion sites

After confirming that disruptions in *E. coli idnK, cpxA*, and *rcsC* were acquired during endosymbiosis, we next investigated the potential mechanisms by which they increase endosymbiont stability. RcsC is an inner membrane-localized histidine kinase (HK) in the multicomponent Rcs phosphorelay system, a response mechanism to environmental stresses that include outer membrane damage, lipopolysaccharide (LPS) synthesis defects, peptidoglycan perturbations, and lipoprotein mislocalization (Figure 2B).^17, 18^ In the Rcs phosphorelay system, extracellular stimuli are received by outer membrane lipoprotein RcsF, which transduces a signal across the periplasm to IgaA, an inhibitor of RcsC.^19^ Activated IgaA derepresses RcsC, leading to RcsC-mediated autophosphorylation of H479, followed by a series of phosphotransfers to D875, and then ultimately to RcsB.^20^ RcsA partners with RcsB to upregulate CPS-mediated capsule biosynthesis, downregulate *flhDC*-mediated flagellum formation and motility, and upregulate additional genes including *rprA, osmC*, and *ftsZ*. RprA is a positive regulator of stress sigma factor RpoS, and a negative regulator of flagellum synthesis. OsmC is a periplasmic peroxidase, and FtsZ is the primary scaffolding protein of cell division. The *rcsC* transposon insertion site observed in EB1 is at amino acid 702 producing a truncated RcsC variant containing the RcsC periplasmic domain, transmembrane domains, and HK domain. A deletion strain of *rcsC, ΔrcsC*::spec^R^, was unable to confer the EB1 phenotype suggesting that the truncated protein is functional.

Similar to RcsC, CpxA is an inner membrane-localized HK that in conjunction with CpxR forms the CpxAR two-component signal transduction system which senses environmental stress including defects in inner membrane protein secretion, misfolding of inner membrane or periplasmic proteins, and lipoprotein export defects (Figure 2C). CpxP derepresses CpxA leading to autophosphorylation of CpxA H248. This phosphate is then transferred to D51 of the cytoplasmic response regulator CpxR, which once phosphorylated binds DNA and activates a transcriptional program. Gene activation affects pili expression and cell adhesion, regulation of secretion systems involved in virulence, efflux of modified peptidoglycan, and metal ion redox homeostasis conferring resistance to antibiotics.^21, 22^ The *cpxA* transposon insertion site in EB1 is positioned at amino acid 143 of the periplasmic domain, hence CpxA-mediated cytoplasmic signaling appears to be abolished entirely. It is not clear why the Tn insertion site in *cpxA* appears to be conserved.

IdnK is a thermosensitive D-gluconate kinase that catalyzes the conversion of D-gluconate and ATP to 6-phos-pho-D-gluconate, a core metabolite of the pentose phosphate pathway and glycolysis (Figure 2D).^7^ The *idnk* transposon insertion site in EB1 is positioned at amino acid 167 (of 187), which based on an alpha-fold model truncates an alpha helix that extends from the globular structure at the C-terminus. Mutants in ED dehydratase (gene *edd*), which catalyzes the step after IdnK in the Entner-Doudoroff pathway,^23^ are defective in flagellum biosynthesis (*flhDC* operon), an operon regulated by RcsC. *E. coli* defective in *edd* and *flhDC* display 15-30% increase in growth rate on media containing sugar as a sole carbon source. Again, it is not clear why the Tn insertion site appears to be conserved or how it may modulate or abolish protein activity.

### EB1 Displays Superior Growth in High Osmotic Pressure

One potential mechanism by which altered RcsC and CpxA signaling may affect endosymbiont stability is by modulating growth rates of bacteria in the high osmolarity of the yeast cytosol. To investigate this possibility, a set of *ΔnadA* mutant *E. coli* strains were characterized in bacterial growth assays with and without 1 M sorbitol, a concentration comparable to the osmotic pressure inside yeast (Figure 3A and 3B).^24^ The parent strain *ΔnadA*, and mutant strains *cpxA*::Tn and *idnK*::Tn exhibit a similar growth rate in media with and without sorbitol, indicating that neither *cpxA* or *idnK* likely affect bacterial growth rate in yeast. In contrast, strain EB1 exhibits a 2.5-fold increase in maximal growth rate in sorbitol containing media relative to the parent *ΔnadA* strain. A comparable 2-3 fold maximal growth rate was observed for *rcsC*::Tn, Δ*rcsC*, as well as a *ΔnadA* strains in which the RcsC phosphorylation sites at 479 and 875 were individually mutated to glutamine. These findings were replicated in sucrose (0.5 M) media, but not in media devoid of osmoprotectant indicating an osmolarity-induced RcsC-dependent growth phenotype. We hypothesize that this mutant RcsC dependent change in growth rate may contribute to the ability of the bacterium to survive in the high osmolarity of the yeast cytosol.

**Figure 3.**
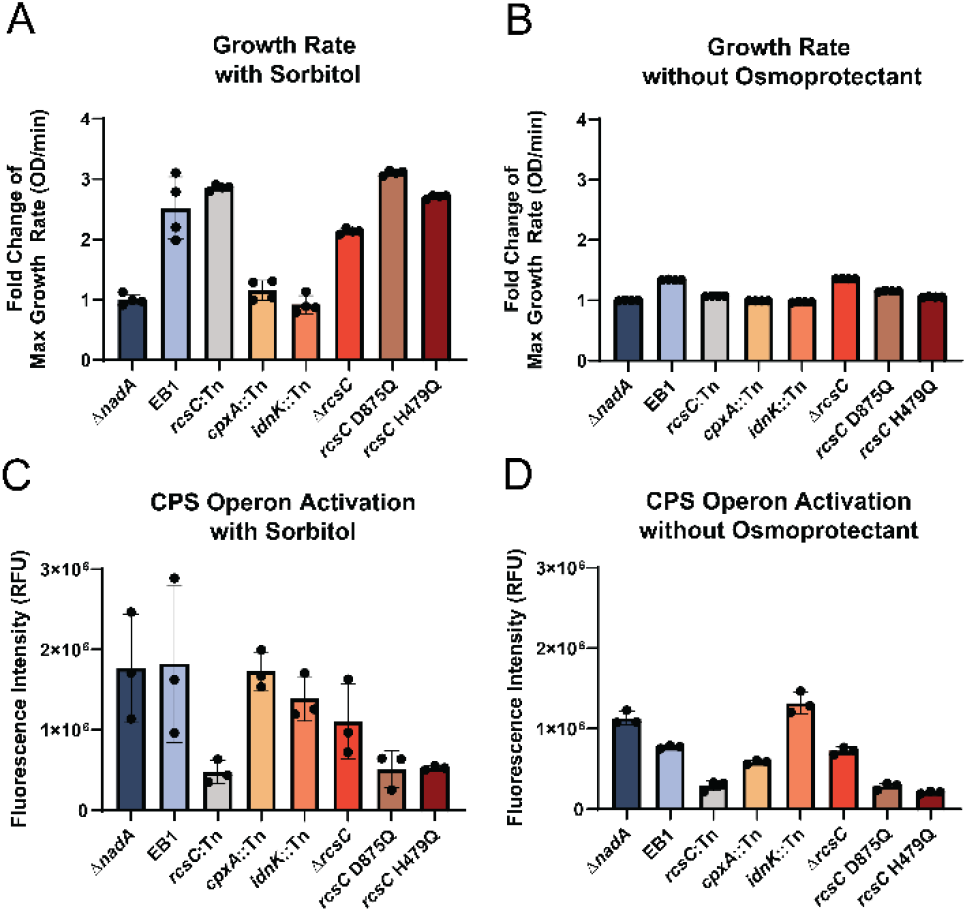
(A) Bacterial growth assays of mutant *E. coli* strains with sorbitol (1 M) and (B) without sorbitol. Values are provided as a fold change of the maximum growth rate relative to Δ*nadA*. (C) Activation of CPS pathway transcription is quantified from the fluorescence of DsRed Express II for each mutant *E. coli* strain with sorbitol and (D) without sorbitol. Each data value is an average of three replicates with error bars representing the standard deviation.

### EB1 Recovers Deleterious Effects on Capsule Biosynthesis

Another possible mechanism by which altered RcsC and CpxA signaling might affect endosymbiont coexistence is by increasing colanic acid biosynthesis and capsule formation as an additional protective strategy beyond expression of the Snare-like proteins. Indeed, upregulation of CPS-mediated capsule biosynthesis is a known down-stream effect of RcsC signaling. Unfortunately, initial attempts to stain the capsule in the endosymbiont produced ambiguous results.^25^ Therefore, a CPS pathway reporter was constructed by assembling a plasmid that encodes the promoter sequence of the CPS operon upstream of a gene encoding fluorescent protein, *dsRed* Express II (Figure S1). Surprisingly, *rcsC*::Tn, H479Q, and D875Q mutations in RcsC caused a reduction in CPS operon expression compared to *ΔnadA* that is independent of osmotic pressure (Figure 3C and 3D). However, for *ΔrcsC* there is an increase in CPS operon expression relative to the mutant strains, indicating that a secondary activation mechanism of CPS exists in the absence of RcsC. We found that both EB1 and *cpxA*::Tn produced a CPS expression profile similar to *ΔnadA E. coli*; all three strains appears to drive CPS expression in an osmoprotectant-dependent manner. These results suggest that while the direct impact of *rcsC*::Tn in EB1 is CPS repression, this effect is offset by an indirect CPS activation mechanism that restores protective CPS activation in high pressure environments, possibly mediated by the *cpxA*::Tn mutation.

### Transcriptomics Analysis of EB1 and *rcsC*::Tn

To evaluate the change in global gene expression within EB1, we prepared log phase and stationary phase cultures of EB1, *rcsC*::Tn and Δ*nadA* in endosymbiont growth medium without antibiotic and performed transcriptomics. In stationary phase cells, we observe an abundance of up- and down-regulated genes in EB1 relative to Δ*nadA*, while in *rcsC*::Tn gene expression is primarily down-regulated relative to Δ*nadA* (Figure 4A and 4B). Interestingly, the primary up-regulated pathways in EB1 center on the degradation of 6-membered ring metabolites (aromatic and pyranose) which likely increase intracellular ATP levels that impact bacterial and yeast growth and provide an alternative carbon source. Notably, none of the top five dysregulated gene pathways in EB1 relative to Δ*nadA* (Enterobactin biosynthesis, the pathway converting trans-cinnamate or 3-hydroxycinnamate to 2-hydroxypentadienoate, upregulation of *lacZ* (LacZ converts lactose to glucose), the *cynS* pathway for cyanate degradation, and *sriD*-mediated sorbitol degradation) were among the top 25 dysregulated gene pathways in *rcsC*::Tn (Figure 4C and 4D).^26^ In *rcsC*::Tn the most affected pathway is the down-regulation of colanic acid synthesis, a segment of the CPS operon. This finding is in agreement with the CPS assay described above (Figure 3C), which showed that the CPS operon is significantly downregulated in *rcsC*::Tn, but not in EB1 where it is likely compensated by transposition insertion in *cpxA*. The collective effects of transposon-disrupted *cpxA, idnK*, and *rcsC* produce a unique gene expression profile in EB1 that promotes endosymbiont longevity, which may explain why the transposons are acquired at the same time (Figure 2).

**Figure 4.**
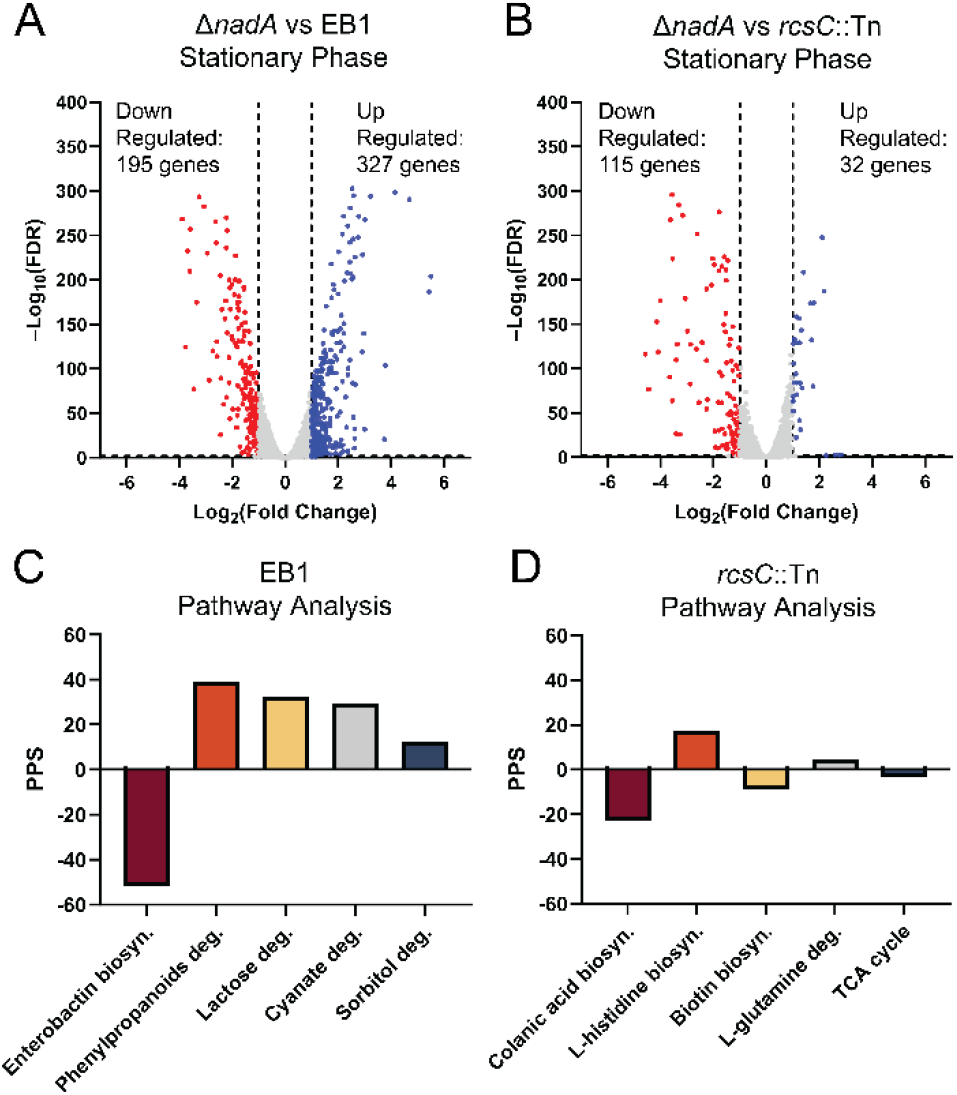
Transcriptomics analysis of EB1 and *rcsC*::Tn compared to Δ*nadA*. (A) Volcano plots depicting up- (blue) and down-regulated (red) genes that experience >2-fold change in expression for Δ*nadA* vs EB1 and (B) Δ*nadA* vs *rcsC*::Tn in stationary phase growth. (C) The top five pathways ranked by pathway perturbation score (PPS) are shown for EB1 and (D) *rcsC*::Tn.

**Figure 5.**
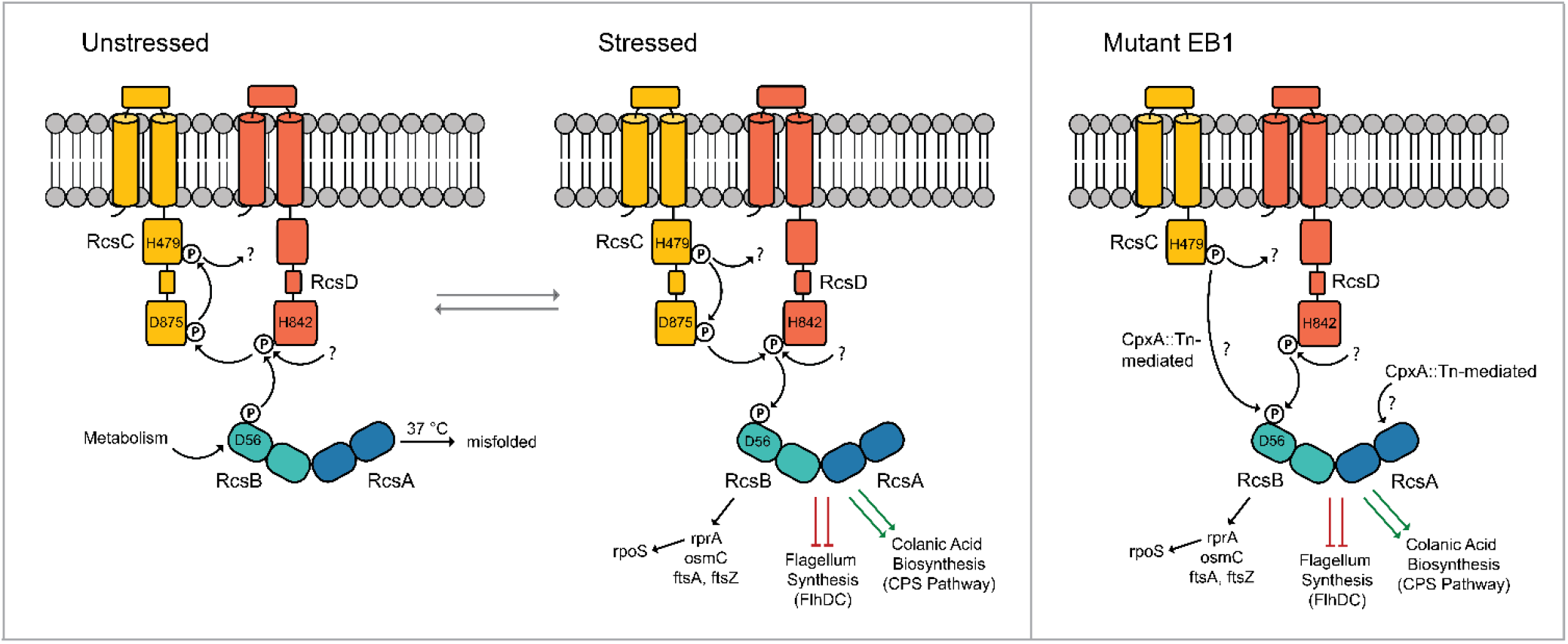
Proposed signaling of the Rcs phosphorelay system in unstressed conditions, conditions that impart stress to the exterior cell envelope, and in EB1 where an IS4-like element has inserted into the protein at amino acid 702. EB1 Rcs signaling indicates cross-talk between the disrupted activities of CpxA and IdnK.

## CONCLUDING REMARKS

In summary, we employed a model system to understand the early genetic alterations that a bacterium engulfed by a higher order host species such as a yeast or by extrapolation an archaeon may have experienced during the early stages of mitochondrial endosymbiont evolution. We have documented that three genes *idnK, cpxA*, and *rcsC* acquire transposon insertions reproducibly at the same position and approximate passage number. The driver for improved endosymbiont phenotype appears to be the *rcsC* transposon insertion, which alone is sufficient to provide improved bacterial growth rates in high osmotic pressure. While this mutation appears to be the driving force for the EB1 phenotype in our fusion experiments and bacterial growth assays, the capsule biosynthesis reporter and transcriptomics analysis revealed a far more complex genetic alteration landscape.

The collective evidence suggests that this transposon insertion comes with disadvantageous effects (i.e., loss of capsule biosynthesis), which require compensating mutations in additional genes. (Figure 4). Our study suggests that transposon insertions in *idnK* and *cpxA* restore a potentially undesirable reduction in capsule biosynthesis and compensate for the downregulation of gene expression that we observe for *rcsC*::Tn by upregulating a series of *rcsC* independent biosynthetic pathways. This is evident in our analysis of the shared regulation of biosynthetic pathways between EB1 and *rcsC*::Tn. Thus the improved stability of EB1 as an endosymbiont is not simply a result of improved growth rate in a high osmotic pressure environment, but the cohesive regulation of a complex transcriptional array. The advantageous genotype that the collective transposons impart on EB1 enable a more stable endosymbiont to form and survive, which is documented by fluorescent microscopy. Although the above study highlights the need to overcome growth rate disparities and environmental stress when two distinct organisms enter in a symbiotic relationship, further analysis is required to elucidate the mechanistic details of this novel bacterial stress response mediated adaption mechanism and the interplay of the associated genes.

We believe that there are several opportunities to improve our model endosymbiont to better recapitulate endosymbiosis and mitochondrial evolution. First, a design with a bacterial-yeast co-dependence that does not depend on auxotrophy in an essential bacterial cofactor or metabolite (i.e. NAD+) would remove genetic alterations induced by metabolic auxotroph starvation.^27^ Second, a strategy that strictly prevents bacterial growth outside of yeast independent of media. Third, further naturally selected (Tn IS) genomic minimization in the endosymbiont, and removal of the bacterial cell wall to reduce genetic alterations mediated by non-mitochondrial components of the bacterium.^6, 7^ To facilitate the removal of the bacterial cell wall, we are inspired by the studies of Errington and coworkers who have engineered synthetic bacterial cell-wall-less L-forms of *E. coli* and *Bacillus subtilis*, which also form and survive naturally within macrophages.^28, 29^

The endosymbiont platform described here has proven its utility as a novel model system to study the early stages of mitochondrial formation by recapitulating bacterial engulfment by a high-order host and monitoring spontaneous mutations that arise as a result of enforced codependence. In addition to their utility as models for the study of early evolutionary events, these endosymbionts may provide unexplored platforms for biosynthesis that benefit from the advantages of each independent organism. Improving the stability of these systems will increase their broad adoption for bioengineering and synthetic biology applications.

## METHODS

### *Escherichia coli* and *Saccharomyces cerevisiae* Strains

All *E. coli* strains used in this study were derived from *E. coli* DH10B strain (F– *mcrA* Δ(*mrr*-*hsdRMS*-*mcrBC*) Φ80*lacZ*Δ*M15* Δ*lacX74 recA1 endA1 araD139* Δ(*ara leu*) 7697 *galU galK rpsL nupG* Δ*nadA*::*gfp*-*kanR* λ– Δ). Construction of DH10B Δ*nadA*::*gfp*-*kanR* was described previously.^9^ Prior to yeast fusions, *E. coli* cells were transformed with plasmid pAM136 encoding an ADP/ATP translocase (*ntt1*) and three SNARE-like proteins (Ctr-incA, Cca_incA, and CT_813).^9^ S. cerevisiae ρ+ NB97 yeast strain (MATa leu2-3,112 lys2 ura3-52 his3ΔHindIII arg8Δ::URA3 [cox2-60::ARG8m] was used for all endosymbiont experiments.^30^

*E. coli* were routinely cultured in 2x yeast extract tryptone (YT) or lysogeny broth (LB). Where indicated, media was supplemented with 50 µg/L kanamycin, 50 µg/L chloramphenicol, 50 µg/L carbenicillin, 50 µg/L spectinomycin or 1 mM L-arabinose. Yeast strains were routinely cultured in YPD medium (1% Bacto yeast extract, 2% Bacto peptone, 2% glucose). Where indicated, 2% glucose was substituted with 0.1% glucose/3% glycerol or 3% glycerol (YPSDG). Sorbitol (1 M) or sucrose (0.5 M) were added to increase osmolality of media.

### *Escherichia coli*-*S. cerevisiae* Endosymbiont Formation

Synthetic endosymbionts were generated and cultured as described previously.^9^ Bacteria were recovered from endosymbionts as described previously.

### Wide-field Fluorescent Microscopy of Endosymbionts

Endosymbionts were grown on YPSGD plates supplemented with 0.1% carbenicillin at 25 °C for 72 hours. The cells were scrapped from the plates and collected by centrifugation (3 min at 845 x *g*) and washed with phosphate buffered saline (PBS). The cell pellets were resuspended in 10 µl of PBS and 3 µl of cell suspension was transferred to a microscope slide and coverslipped. Images of endosymbionts were acquired using a widefield epifluorescent Nikon Ti2-E inverted microscope (Ex. 466 nm, Em. 525 nm).

### Whole-genome Sequencing of Evolved Bacteria

EB1 cells were cultured overnight in LB supplemented with kanamycin and chloramphenicol (pAM136 and genomic selection markers), pelleted, and the genome extracted and analyzed by whole genome sequencing (Next Generation Sequencing (NGS), GENEWIZ, Inc.). Transposon insertions were confirmed by PCR amplification and sequencing of *rcsC* (primers ‘rcsC del conf for’+ ‘rcsC del conf rev’), *cpxA* (primers ‘cpxA conf for 2’+ ‘cpxA conf rev’) and *idnK* (primers ‘idnK conf for 2’ + idnK conf rev’)(Table S1). Three primer pairs were used to monitor transposons over-time in follow-up experiments.

### Construction of *E. coli* KO and Genomic Mutant Strains

All primers used for the generation and characterization of *E. coli* KO strains are listed in Table S1. Gene disruption cassettes were generated by PCR. Gene disruption cassettes were introduced into target sites in the *E. coli* genome using Lambda (λ) Red-mediated recombineering.^31^ Detailed description of *E. coli* KO strain generation is summarized in Table S2. The H479Q and D875Q point mutations were introduced into *E. coli* DH10B using a CRISPR-Cas approach.^32^ All primers and gRNA sequences used to generate mutant *E. coli* are provided in Table S1. A detailed protocol is provided in the supporting information.

### Bacterial Growth Assay

Bacterial cultures were inoculated to a starting OD_600_ of 0.0025 in YPSDG with and without sorbitol (1 M) from an overnight culture. Growth curves were collected in quadruplicate using a BioTek Logphase 600 plate reader (Agilent, CA, USA) in 96 well plates in triplicate. Fold change of max growth rate values were determined by the instrument for each growth assay.

### CPS-Reporter Assay

To assemble the CPS operon reporter plasmid pDD286 (Figure S1), the 459 base pair region^33^ immediately upstream of the *E. coli wza* gene were inserted in front of a red fluorescence protein dsRedExpress II on a plasmid backbone encoding a ColE-1 origin and spec^R^. Each *E. coli* strain was transformed with pDD286. Bacterial cultures were inoculated to a starting OD_600_ of 0.0025 in YPSDG with and without sorbitol (1 M) from an overnight culture. The fluorescence (Ex. 550 nm, Em. 590 nm) of each culture was measured using a SpectraMax iD3 fluorescence plate reader in 96 well plates in triplicate.

### Transcriptomics of *E. coli* EB1, Δ*nadA*, and *rcsC*::Tn

Bacterial cultures were inoculated (OD_600_= 0.0025) in YPSDG and grown to either OD_600_: 0.6 (log phase) or overnight (stationary phase). Three 1 mL replicates of each sample were pelleted, frozen, and the transcriptome was analyzed (GENEWIZ, Inc.). The gene expression fold change of EB1 and *rcsC*::Tn was determined relative to Δ*nadA*. Gene expression pathway sorting was performed using EcoCyc.^26^

## ASSOCIATED CONTENT

Experimental methods, nucleotide sequences, a supplementary figure and tables.

## Author Contributions

L.S., D.A.D., and P.G.S. were responsible for the study conception and experimental design. L.S., H. Z., B. B. L., I.H.B., C. N. and D.A.D. performed the experiments. L.S., D.A.D., and P.G.S. analyzed data and wrote the manuscript.

## Funding Sources

We acknowledge Kristen Williams for her assistance in manuscript preparation. This work was supported by NIH Grant RGM145323A to P.G.S.

## ACKNOWLEDGMENT

*We thank Frantisek Supek and Christian Diercks for helpful discussions and Kathryn Spencer for help with widefield microscopy*

## ABBREVIATIONS

S. cerevisiae: Saccharomyces cerevisiae.
E. coli: Escherichia coli
COX2: cytochrome c oxidase subunit 2
NAD+: Nicotanimide
EB1: Evolved Bacteria 1
Tn: transposon
IS: insertion sequences
HK: Histidine Kindase
LPS: lipopolysaccharide
ED: Entner-Dou-doroff
KO: knockout
gfp: green fluorescent protein

## Notes

### Competing Interest Statement

The authors have declared no competing interest.

## REFERENCES

(1) Sagan, L. On the origin of mitosing cells. J. Theor. Biol. 1967, 14 (3), 255–274. DOI: 10.1016/0022-5193(67)90079-3.

(2) Gray, M. W. Lynn Margulis and the endosymbiont hypothesis: 50 years later. Mol. Biol. Cell 2017, 28 (10), 1285–1287. DOI: 10.1091/mbc.E16-07-0509.

(3) John, P.; Whatley, F. R. Paracoccus denitrificans and the evolutionary origin of the mitochondrion. Nature 1975, 254 (5500), 495–498. DOI: 10.1038/254495a0.

(4) Andersson, S. G.; Zomorodipour, A.; Andersson, J. O.; Sicheritz-Ponten, T.; Alsmark, U. C.; Podowski, R. M.; Naslund, A. K.; Eriksson, A. S.; Winkler, H. H.; Kurland, C. G. The genome sequence of Rickettsia prowazekii and the origin of mitochondria. Nature 1998, 396 (6707), 133–140. DOI: 10.1038/24094.

(5) Geiger, O.; Sanchez-Flores, A.; Padilla-Gomez, J.; Degli Esposti, M. Multiple approaches of cellular metabolism define the bacterial ancestry of mitochondria. Sci. Adv. 2023, 9 (32), eadh0066. DOI: 10.1126/sciadv.adh0066.

(6) Roger, A. J.; Munoz-Gomez, S. A.; Kamikawa, R. The Origin and Diversification of Mitochondria. Curr. Biol. 2017, 27 (21), R1177– R1192. DOI: 10.1016/j.cub.2017.09.015.

(7) Peekhaus, N.; Conway, T. What’s for dinner?: Entner-Doudoroff metabolism in Escherichia coli. J. Bacteriol. 1998, 180 (14), 3495–3502. DOI: 10.1128/JB.180.14.3495-3502.1998.

(8) Zimmer, C. Origins. On the origin of eukaryotes. Science 2009, 325 (5941), 666–668. DOI: 10.1126/science.325_666.

(9) Mehta, A. P.; Supekova, L.; Chen, J. H.; Pestonjamasp, K.; Webster, P.; Ko, Y.; Henderson, S. C.; McDermott, G.; Supek, F.; Schultz, P. G. Engineering yeast endosymbionts as a step toward the evolution of mitochondria. Proc. Natl. Acad. Sci. U. S. A. 2018, 115 (46), 11796–11801. DOI: 10.1073/pnas.1813143115.

(10) Schmitz-Esser, S.; Linka, N.; Collingro, A.; Beier, C. L.; Neuhaus, H. E.; Wagner, M.; Horn, M. ATP/ADP translocases: a common feature of obligate intracellular amoebal symbionts related to Chlamydiae and Rickettsiae. J. Bacteriol. 2004, 186 (3), 683–691. DOI: 10.1128/JB.186.3.683-691.2004 From NLM Medline.

(11) Supekova, L.; Supek, F.; Greer, J. E.; Schultz, P. G. A single mutation in the first transmembrane domain of yeast COX2 enables its allotopic expression. Proc. Natl. Acad. Sci. U. S. A. 2010, 107 (11), 5047–5052. DOI: 10.1073/pnas.1000735107.

(12) Delevoye, C.; Nilges, M.; Dehoux, P.; Paumet, F.; Perrinet, S.; Dautry-Varsat, A.; Subtil, A. SNARE protein mimicry by an intracellular bacterium. PLoS Pathog. 2008, 4 (3), e1000022. DOI: 10.1371/journal.ppat.1000022.

(13) Sulo, P.; Griac, P.; Klobucnikova, V.; Kovac, L. A method for the efficient transfer of isolated mitochondria into yeast protoplasts. Curr. Genet. 1989, 15 (1), 1–6. DOI: 10.1007/BF00445745.

(14) Mehta, A. P.; Ko, Y.; Supekova, L.; Pestonjamasp, K.; Li, J.; Schultz, P. G. Toward a Synthetic Yeast Endosymbiont with a Minimal Genome. J. Am. Chem. Soc. 2019, 141 (35), 13799–13802. DOI: 10.1021/jacs.9b08290.

(15) Consuegra, J.; Gaffe, J.; Lenski, R. E.; Hindre, T.; Barrick, J. E.; Tenaillon, O.; Schneider, D. Insertion-sequence-mediated mutations both promote and constrain evolvability during a long-term experiment with bacteria. Nat. Commun. 2021, 12 (1), 980. DOI: 10.1038/s41467-021-21210-7 From NLM Medline.

(16) Cesar, S.; Anjur-Dietrich, M.; Yu, B.; Li, E.; Rojas, E.; Neff, N.; Cooper, T. F.; Huang, K. C. Bacterial Evolution in High-Osmolarity Environments. mBio 2020, 11 (4). DOI: 10.1128/mBio.01191-20.

(17) Sato, T.; Takano, A.; Hori, N.; Izawa, T.; Eda, T.; Sato, K.; Umekawa, M.; Miyagawa, H.; Matsumoto, K.; Muramatsu-Fujishiro, A.; et al. Role of the inner-membrane histidine kinase RcsC and outer-membrane lipoprotein RcsF in the activation of the Rcs phosphorelay signal transduction system in Escherichia coli. Microbiology 2017, 163 (7), 1071–1080. DOI: 10.1099/mic.0.000483.

(18) Mitchell, A. M.; Silhavy, T. J. Envelope stress responses: balancing damage repair and toxicity. Nat. Rev. Microbiol. 2019, 17 (7), 417–428. DOI: 10.1038/s41579-019-0199-0.

(19) Delhaye, A.; Collet, J. F.; Laloux, G. A Fly on the Wall: How Stress Response Systems Can Sense and Respond to Damage to Peptidoglycan. Front Cell Infect. Microbiol. 2019, 9, 380. DOI: 10.3389/fcimb.2019.00380.

(20) Wall, E.; Majdalani, N.; Gottesman, S. The Complex Rcs Regulatory Cascade. Annu. Rev. Microbiol. 2018, 72, 111–139. DOI: 10.1146/annurev-micro-090817-062640.

(21) Flores-Kim, J.; Darwin, A. J. Regulation of bacterial virulence gene expression by cell envelope stress responses. Virulence 2014, 5 (8), 835–851. DOI: 10.4161/21505594.2014.965580.

(22) Ruiz, N.; Silhavy, T. J. Sensing external stress: watchdogs of the Escherichia coli cell envelope. Curr Opin Microbiol 2005, 8 (2), 122–126. DOI: 10.1016/j.mib.2005.02.013.

(23) Leatham, M. P.; Stevenson, S. J.; Gauger, E. J.; Krogfelt, K. A.; Lins, J. J.; Haddock, T. L.; Autieri, S. M.; Conway, T.; Cohen, P. S. Mouse intestine selects nonmotile flhDC mutants of Escherichia coli MG1655 with increased colonizing ability and better utilization of carbon sources. Infect. Immun. 2005, 73 (12), 8039–8049. DOI: 10.1128/IAI.73.12.8039-8049.2005.

(24) Hutchison, H. T.; Hartwell, L. H. Macromolecule synthesis in yeast spheroplasts. J. Bacteriol. 1967, 94 (5), 1697–1705. DOI: 10.1128/jb.94.5.1697-1705.1967 From NLM Medline.

(25) Breakwell, D. P.; Moyes, R. B.; Reynolds, J. Differential staining of bacteria: capsule stain. Curr. Protoc. Microbiol. 2009, Appendix 3, Appendix 3I. DOI: 10.1002/9780471729259.mca03is15.

(26) Karp, P. D.; Paley, S.; Caspi, R.; Kothari, A.; Krummenacker, M.; Midford, P. E.; Moore, L. R.; Subhraveti, P.; Gama-Castro, S.; Tierrafria, V. H.; et al. The EcoCyc Database (2023). EcoSal Plus 2023, 11 (1), eesp00022023. DOI: 10.1128/ecosalplus.esp-0002-2023.

(27) Chen, Y.; Ying, Y.; Lalsiamthara, J.; Zhao, Y.; Imani, S.; Li, X.; Liu, S.; Wang, Q. From bacteria to biomedicine: Developing therapies exploiting NAD(+) metabolism. Bioorg. Chem. 2024, 142, 106974. DOI: 10.1016/j.bioorg.2023.106974.

(28) Mercier, R.; Kawai, Y.; Errington, J. Excess membrane synthesis drives a primitive mode of cell proliferation. Cell 2013, 152 (5), 997–1007. DOI: 10.1016/j.cell.2013.01.043.

(29) Kawai, Y.; Mickiewicz, K.; Errington, J. Lysozyme Counteracts beta-Lactam Antibiotics by Promoting the Emergence of L-Form Bacteria. Cell 2018, 172 (5), 1038–1049 e1010. DOI: 10.1016/j.cell.2018.01.021 From NLM Medline.

(30) Bonnefoy, N.; Fox, T. D. Genetic transformation of Saccharomyces cerevisiae mitochondria. Methods Cell Biol. 2001, 65, 381–396. DOI: 10.1016/s0091-679x(01)65022-2 From NLM Medline.

(31) Datsenko, K. A.; Wanner, B. L. One-step inactivation of chromosomal genes in Escherichia coli K-12 using PCR products. Proc. Natl. Acad. Sci. U. S. A. 2000, 97 (12), 6640–6645. DOI: 10.1073/pnas.120163297 From NLM Medline.

(32) Jiang, W.; Bikard, D.; Cox, D.; Zhang, F.; Marraffini, L. A. RNA-guided editing of bacterial genomes using CRISPR-Cas systems. Nat. Biotechnol. 2013, 31 (3), 233–239. DOI: 10.1038/nbt.2508.

(33) Rahn, A.; Whitfield, C. Transcriptional organization and regulation of the Escherichia coli K30 group 1 capsule biosynthesis (cps) gene cluster. Mol. Microbiol. 2003, 47 (4), 1045–1060. DOI: 10.1046/j.1365-2958.2003.03354.x.

