## Supplementary Information for "Natural Selection in a Synthetic Yeast Endosymbiont Promotes Stable Coexistence"

**This PDF file includes:**

Figure S1
Tables S1 and S2

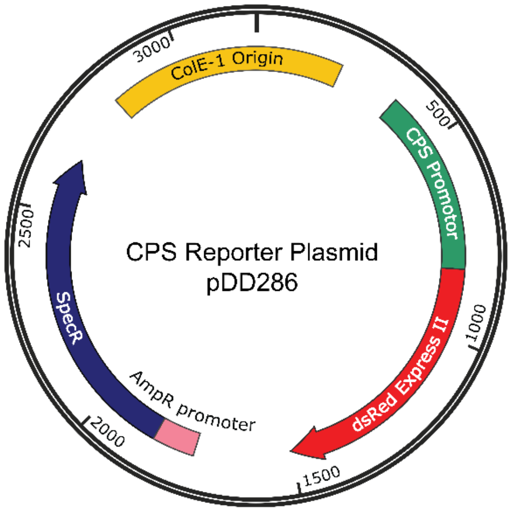


**Figure S1.** Plasmid map of CPS reporter plasmid pDD286. The plasmid encodes the 459 bp sequence upstream of *E. coli wza* as a transcriptional element for measuring CPS pathway expression, which controls capsule biosynthesis. Activation of transcription produces dsRED Express II, a red fluorescent protein. The 3186 bp plasmid encodes a spectinomycin resistance gene (spec$^R$) and a ColE-1 bacterial origin of replication. Color coded sequence of the promoter and fluorescent protein is provided in Table S1 (sequence oDAD1006).

Construction of *E. coli* DH10B *rcsC*::H479Q and *rcsC*::D875Q strains

H(CAT)479Q(CAG) point mutation was introduced into the *E. coli* DH10B genome using a CRISPR-Cas approach. All oligonucleotides used for generation of the mutant *E. coli* strain are listed in Table S1. DH10B cells were transformed with CAS9BAC1P plasmid (Sigma) and maintained at 30 °C in LB with kanamycin. CRISPR30:LacZ plasmid (Sigma, amp$^R$) was mutagenized by Q5 site-directed mutagenesis kit (NEB) and primers 'rcsC H479 for' and 'rcsC H479 rev' to yield a plasmid that encodes gRNA targeting the wild type *rcsC* gene at position H479. The sequence was verified for the resulting plasmid CRISP30:RcsC-H479 with 'J23119 for4' and 'J23119 rev4' primers. In the next step, we generated a DNA fragment of the *rcsC* gene that contained the H479 mutation. Two pairs of primers 'rcsC H479Q up for' and 'rcsC H479Q up rev', and 'rcsC H479Q down for' and 'rcsC H479Q down rev' were used to amplify two *rcsC* gene fragment from DH10B genomic DNA. Two resulting fragments were then used as the templates in a fusion PCR reaction using primers 'rcsC H479Q for' and 'rcsC H479Q rev' to yield 'rcsC H479Q template' (1 kb with H479Q mutation in the middle of the PCR fragment). DH10B (CAS9BAC1P) cells were co-transformed with the CRISPR:RcsC-H479 and the 'rcsC H479Q template' DNA and transformants were selected on LB supplemented with kanamycin and ampicillin at 30 °C. Briefly to generate *rcsC* D(GAC)875Q(CAG), CRISPR30:LacZ plasmid (Sigma, amp$^R$) was mutagenized by Q5 site-directed mutagenesis kit and primers 'oDAD996' and 'oDAD997' to yield a plasmid that encodes gRNA targeting the *rcsC* gene sequence at D875. A dsDNA doner DNA fragment was ordered as a gene block 'oDAD995' (IDT). The presence of the *rcsC* H479Q and D875Q mutations were identified in transformants by amplifying the *rcsC* gene from chromosomal DNA with 'oDAD1996' and 'oDAD1997' and DNA sequencing.

**Table S1**

PCR oligonucleotides used for construction of gene disruption cassettes and plasmids.

| **Oligonucleotide name** | **Oligonucleotide sequence** |
|---|---|
| rcsC del conf for (51) | ATTATCGCTGACCGCTTTGT |
| rcsC del conf rev (52) | TGGTTTGGCTATACGGAAGG |

| | |
|---|---|
| cpxA conf for 2 (LS60) | AGATGGTCACCCGTGGTTTA |
| cpxA conf rev (LS59) | ATCAGGCATCCTGCTCAAAT |
| idnK conf for 2 (LS60) | TCGCAGCAGGTAAGATGATTCA |
| idnK conf rev (LS56) | ACCGAACTGTTCCCGATGT |
| rcsC-specR del for | GTTAGCAGGATCGTCGCTCTCCATCTCGATCAAATAAATACGCGCCAACATTATTTGCCGACTACCTTGGTGATCTCGCC |
| rcsC-specR del rev | AAGCATCGTGTATTCGGTGCCGGTTGATAAGGTGCTGGAACGCATTCGCACCGCCGCAGTCTCACGCCCGG |
| spec for | GAACATAGCGTTGCCTTGGT |
| spec rev | ATGTCATTGCGCTGCCATTC |
| rcsD-specR del for | CCCTATCACTTCGCGAAGTTTTAACAGGTCATAAACACGATTATTTGCCGACTACCTTGGTGATCTCGCC |
| rcsD-specR del rev | AATAATTACGTTCATATTGTTCATGTAATAGGCTACCTTGCCGCCGCAGTCTCACGCCCGG |
| rcsD del conf for | AACAAAACCTTCACTCGCAAC |
| rcsD del conf rev | ATGCCATCGCCGTACTTATC |
| FRT PGK for | GGGGGGGGGGGGCGGCCGCGAAGT |
| FRT PGK rev | GACTCACTATAGGGCTCGAGGAA |
| rcsC-EB1 for1 | TGCGTCGCTCTGTCAGTTC |
| rcsC-EB1 rev1 | CCCCCCCCCCCCGCTCGTGGCTGCAACCCTACTA |
| rcsC-EB1 for2 | GAACTTCCTCGAGCCCTATAGTGAGTCAGATCAGTTGGGATCGTTGG |
| rcsC-EB1 rev2 | CCCTCTCGGCATATAACGTC |
| rcsC-EB1 final for1 | AGGGCAGAGCGGTAGTGAC |
| rcsC-EB1 final rev1 | GTCAAGCGGTAACCATCCAT |
| rcsC conf for (LS101) | CCCTCTCGGCATATAACGTC |
| rcsC conf rev (LS78 | TGCGTCGCTCTGTCAGTTC |
| kan for (LS42) | GGATGATCTGGACGAAGAGC |
| kan rev (LS69) | CGAGACTAGTGAGACGTGCT |
| nadA-gfp-kan for (LS5) | CGAGATGGTAAGATGAGCGTAA |
| nadA-gfp-kan rev (LS6) | CGCCTTATTATTCGTTATCCAC |
| nadA-gfp-kan confirm for (LS1) | TTCCCCACCATTCGACTATC |
| nadA-gfp-kan confirm rev (LS2) | CACGTAGTGTAGCCGCAAAA |
| cpxA-EB1 for1 | CGCTGGTGTTGATGTTGGTT |
| cpxA-EB1 rev1 | CCCCCCCCCCCCGCTGTCGTTCTCAAAATCG |
| cpxA-EB1 for2 | GAACTTCCTCGAGCCCTATAGTGAGTCTTTGACCGCCCGCTATTACT |
| cpxA-EB1 rev2 | CGCTCCAGTTCCTTGCTTTC |
| cpxA-EB1 final for1 (LS91) | CGATCATCCGCAGAAGAAA |
| cpxA-EB1 final rev1 (LS92) | ACTCCAGGCCAACCACAAC |
| cpxA conf for | TCGTAAACTGCCGGATCGTA |
| idnK-EB1 for1 (LS76) | TAGCCCCCATGTTCATTTCC |
| idnK-EB1 rev1 | CCCCCCCCCCCCATTAGGGGATTCATCAGAGCA |

| | |
|---|---|
| idnK-EB1 for2 | GAACTTCCTCGAGCCCTATAGTGAGTCCTGTGCTGGCGATACGA C |
| idnK-EB1 final for1 | TAGCCCCCATGTTCATTTCC |
| idnK conf for (LS76) | TAGCCCCCATGTTCATTTCC |
| rcsC-specR 356 for | GTATTTTCATTCCGGCGGAAAGCGACGCCCTGCGACTGGAAGAA CATGAGTGATTATTTGCCGACTACCTTGGTGATCTCGCC |
| rcsC-specR 471 for | CGTTGCAGGAGATGGCACAAGCAGCGGAACAGGCGAGCCAGTC AAAATCGTGATTATTTGCCGACTACCTTGGTGATCTCGCC |
| rcsC-specR 705 for | CCGTTGTACGGCGCTCAGTACCCGCAGAAAAAAGGCGTGGAAGG GTTGAGTGATTATTTGCCGACTACCTTGGTGATCTCGCC |
| rcsC-specR 820 for | CTGCTAACGCTCTGCCGTCAACGGACAAAGCGGTCAGCGATAAT GACGATTGATTATTTGCCGACTACCTTGGTGATCTCGCC |
| rcsC trunc rev | ATCTGGCATTTGCACTGAATGCCGGATGCGGCGTAAACGCCTTAT CCGTCCCGCCGCAGTCTCACGCCCGG |
| rcsC trunc conf for | AGTGAAGTTCTGCGCCTGTT |
| rcsC trunc conf rev | TACTCTCGCACGGATGTACG |
| rcsC H479 for | TGACTGACGGGTTTTAGAGCTAGAAATAGC |
| rcsC H479 rev | TGAGCTGCGAGCTAGCATTATACCTAGG |
| J23119 for4 | TTGAATACTCATACTCTTCCTTTTTCA |
| J23119 rev4 | GATATCAGCGCTTTAAATTTGC |
| rcsC H479Q down for | GCGACGGTAAGCCAGGAATTGCGCAC |
| rcsC H479Q down rev | AACAAGCGCACCACTTGTTT |
| rcsC H479Q up for | GCATTCGCATGTTGATCCTT |
| rcsC H479Q up rev | CGTGGGCAATTCCTGGCTTACCGTCGC |
| rcsC H479Q for | GGTATCGCGAACACGGATAG |
| rcsC H479Q rev | TACTCTCGCACGGATGTACG |
| oDAD979 – add gRNA rev RcsC D875Q by KLD | ACGCATTCGTGCTAGCATTATACCTAGGACTGAGC |
| oDAD980 – add gRNA fwd RcsC D875Q by KLD | TGCGTCAAGGTTTTAGAGCTAGAAATAGCAAGTTAAAATAAGG |
| oDAD995 – doner dsDNA Sequence for D875Q | TCAGCGTCTGTTTTATCACATCCAGCGTTACCGGCTTCGACAGGC AGCTGTCCATACCGGACTCCAGACACCGCTGCTTCTCTTCAGCC AACGCATTAGCAGTTACTCCGATTACCGGCAACGTCAGTCCCAAC TGACGAATGCGTTGCGTCAATCTATAACCATCCATATTTGGCATG TTGACCTGGCTAAGCACGATATCAATATGATTCTTGCTAAGTACAT TAAGCGCATCGACGCCATCATTCGCGGTTTTACATTGATAGCCCA ACGATCCCAACTGATCTGCCAGCAAACGCCGGTTAATCGGATGA TCATCCACGACCAGAATCATCAT |
| oDAD996 – PCR Mut. Verification for D875Q Rev | TCAGCGTCTGTTTTATCACATCCAG |
| oDAD997 – PCR Mut. Verification for D875Q Fwd | ATGATGATTCTGGTCGTGGATGATCATC |
| oDAD1006 – G-block encoding CPS promoter (green), dsRED Express II | GCCTTTGAGTGAGCTGATACCGCTCGCCGCAGCCGAACGACCGA GCGCGTCAACCTAAAGAAACTCCTAAAAACCATATTGAATGACAC TTAATATAATTCTTAAAAATAGCCAATTACCGAATTGTTATCTTGCC TGCTATTCCGTTAGCTGTAACACTTCCTCCTGCATTATTGGAAAG |

| | |
|---|---|
| (red) and homology regions for addition to spec$^R$/ColE-1 backbone | CCAATATTCAATTACCATAATGTCACCTTATATTCGGAAGTTAATTT<br>CCGTAATAACCCTTTCTCATCAACGACTGCATGCTGAGAGGGTGT<br>GAAAAATTAAACTAAATCAGTGTATTGGTAGCTAAAAAGCCAGGG<br>GCGGTAGCGTGTCTGGATGCCTGAAAGACCAGTCTGAATTATTCT<br>GTCAATAGCCTGCGGATGAAATCAATTTTTTTAGGACTGATGCCA<br>GTTAAATTTTTATTCAATTAACTGCAAGGTAAATGTGCATGGCGAC<br>ATTGCTATTAATAGTGCACAGGATAATTACTCTGCCAAAGTGATAA<br>ATAAACAATGAATTCCACTGAGAACGTCATCAAGCCCTTCATGCG<br>CTTCAAGGTGCACATGGAGGGCTCCGTGAACGGCCACGAGTTCG<br>AGATCGAGGGCGAGGGCGAGGGCAAGCCCTACGAGGGCACCCA<br>GACCGCCAAGCTGCAGGTGACCAAGGGCGGCCCCCTGCCCTTC<br>GCCTGGGACATCCTGTCCCCCCAGTTCCAGTACGGCTCCAAGGT<br>GTACGTGAAGCACCCCGCCGACATCCCCGACTACAAGAAGCTGT<br>CCTTCCCCGAGGGCTTCAAGTGGGAGCGCGTGATGAACTTCGAG<br>GACGGCGGCGTGGTGACCGTGACCCAGGACTCCTCCCTGCAGG<br>ACGGCACCTTCATCTACCACGTGAAGTTCATCGGCGTGAACTTCC<br>CCTCCGACGGCCCCGTAATGCAGAAGAAGACTCTGGGCTGGGA<br>GCCCTCCACCGAGCGCCTGTACCCCCGCGACGGCGTGCTGAAG<br>GGCGAGATCCACAAGGCGCTGAAGCTGAAGGGCGGCGGCCACT<br>ACCTGGTGGAGTTCAAGTCAATCTACATGGCCAAGAAGCCCGTG<br>AAGCTGCCCGGCTACTACTACGTGGACTCCAAGCTGGACATCAC<br>CTCCCACAACGAGGACTACACCGTGGTGGAGCAGTACGAGCGC<br>GCCGAGGCCCGCCACCACCTGTTCCAGTAATTAAGCCAGCCCC |

**Table S2**

Construction of *E. coli* genetic knockout strains

| ***E. coli* KO strain** | **PCR reaction** | **PCR DNA template** | **PCR primers** | **PCR products and *E. coli* strains** |
|---|---|---|---|---|
| **DH10B Δ*nadA* Δ*rcsC::spec*$^R$** | 1 | *spec*$^R$ gene cassette | rcsC-specR del for<br>rcsC-specR del rev | PCR product gel purified, transformed into **DH10B Δ*nadA::gfp-kan*$^R$ (pKD46)** |
| | 2 | genomic DNA from #1 | rcsC del conf for<br>spec for | diagnostic PCR, product sequenced to confirm correct gene disruption |
| | 3 | genomic DNA from #1 | rcsC del conf rev<br>spec rev | diagnostic PCR, product sequenced to confirm correct gene disruption → **DH10B Δ*nadA::gfp-kan*$^R$ Δ*rcsC::spec*$^R$ (pKD46)** |
| | | | | **DH10B Δ*nadA::gfp-kan*$^R$ Δ*rcsC::spec*$^R$ (pKD46)** cultured at 42 °C to remove pKD46 |
| **DH10B Δ*nadA* Δ*rcsD::spec*$^R$** | 4 | *spec*$^R$ gene cassette | rcsD-specR del for<br>rcsD-specR del rev | PCR product gel purified, transformed into **DH10B Δ*nadA::gfp-kan*$^R$ (pKD46)** |
| | 5 | genomic DNA from #4 | rcsD del conf for<br>spec for | diagnostic PCR, product sequenced to confirm correct gene disruption |
| | 6 | genomic DNA from #4 | rcsD del conf rev<br>spec rev | diagnostic PCR, product sequenced to confirm correct |

| | | | | |
|---|---|---|---|---|
| | | | | gene disruption → **DH10B** ***ΔnadA::gfp-kan***$^R$ ***ΔrcsD::spec***$^R$ **(pKD46)** |
| | | | | **DH10B** ***ΔnadA::gfp-kan***$^R$ ***ΔrcsD::spec***$^R$ **(pKD46)** cultured at 42 $^o$C to remove pKD46 |
| **DH10B** ***rcsC::trans*** | 7 | FRT-PGK-gb2-neo-FRT cassette | FRT PGK for<br>FRT PGK rev' | PCR product gel purified |
| | 8 | EB1 genomic DNA | rcsC-EB1 for1<br>rcsC-EB1 rev1 | PCR product gel purified |
| | 9 | PCR products 7+8 | rcsC-EB1 for1<br>FRT PGK rev | PCR product gel purified |
| | 10 | EB1 genomic DNA | rcsC-EB1 for2<br>rcsC-EB1 rev2' | PCR product gel purified |
| | 11 | PCR products 9+10 | rcsC-EB1 final for1<br>rcsC-EB1 final rev1 | PCR product gel purified, transformed into **DH10B (pKD46)** |
| | 12 | genomic DNA from #11 | rcsC conf for<br>kan rev | diagnostic PCR, product sequenced to confirm correct gene disruption |
| | 13 | genomic DNA from #11 | rcsC conf rev<br>kan for | diagnostic PCR, product sequenced to confirm correct gene disruption→ **DH10B** ***ΔrcsC::trans-kan***$^R$ **(pKD46)** |
| | | | | **DH10B** ***ΔrcsC::trans-kan***$^R$ **(pKD46)** cultured at 42 $^o$C to remove pKD46 |
| | | | | **DH10B** ***ΔrcsC::trans-kan***$^R$ was transformed with pCP20 |
| | | | | **DH10B** ***ΔrcsC::trans-kan***$^R$ **(pCP20)** was grown at 42 $^o$C to delete *kan*$^R$ and remove pCP20 (*amp*$^R$) → colonies that were sensitive to ampicillin and kanamycin were selected for further characterization |
| | 14 | genomic DNAs from previous step | rcsC conf for<br>rcsC conf rev | diagnostic PCR, product sequenced to confirm correct *kan*$^R$ gene deletion → **DH10B** ***ΔrcsC::trans*** |
| | 15 | genomic DNA from **DH10B** ***ΔnadA::gfp-kan***$^R$ | nadA-gfp-kan for<br>nadA-gfp-kan for | PCR product gel purified, transformed into **DH10B** ***ΔrcsC::trans*** **(pKD46)** |
| | 16 | genomic DNAs from previous step | nadA-gfp-kan for<br>nadA-gfp-kan for | diagnostic PCR, product sequenced to confirm correct *nadA* gene disruption and insertion of *gfp-kan*$^R$ cassette → **DH10B** ***ΔrcsC::trans*** ***ΔnadA::gfp-kan***$^R$ **(pKD46)** |
| | | | | **DH10B** ***ΔrcsC::trans*** ***ΔnadA::gfp-kan***$^R$ **(pKD46)** |

| | | | | |
|---|---|---|---|---|
| | | | | cultured at 42 $^{o}$C to remove pKD46 |
| **DH10B *cpxA::trans*** | 17 | EB1 genomic DNA | cpxA-EB1 for1<br>cpxA-EB1 rev1 | PCR product gel purified |
| | 18 | PCR products 7+17 | cpxA-EB1 for1<br>FRT PGK rev | PCR product gel purified |
| | 19 | EB1 genomic DNA | cpxA-EB1 for2<br>cpxA-EB1 rev2 | PCR product gel purified |
| | 20 | PCR products 18+19 | cpxA-EB1 final for1<br>cpxA-EB1 final rev1 | PCR product gel purified, transformed into **DH10B (pKD46)** |
| | 21 | genomic DNA from #20 | cpxA conf for<br>kan rev | diagnostic PCR, product sequenced to confirm correct gene disruption |
| | 22 | genomic DNA from #20 | cpxA conf rev<br>kan for | diagnostic PCR, product sequenced to confirm correct gene disruption→ **DH10B ∆*cpxA::trans-kan$^{R}$* (pKD46)** |
| | | | | **DH10B ∆*cpxA::trans-kan$^{R}$* (pKD46)** cultured at 42 $^{o}$C to remove pKD46 |
| | | | | **DH10B ∆*cpxA::trans-kan$^{R}$*** was transformed with pCP20 |
| | | | | **DH10B ∆*cpxA::trans-kan$^{R}$* (pCP20)** was grown at 42 $^{o}$C to delete *kan$^{R}$* and remove pCP20 (*amp$^{R}$*) → colonies that were sensitive to ampicillin and kanamycin were selected for further characterization |
| | 23 | genomic DNAs from previous step | cpxA-EB1 final for1<br>cpxA-EB1 final rev1 | diagnostic PCR, product sequenced to confirm correct *kan$^{R}$* gene deletion → **DH10B ∆*cpxA::trans*** |
| | 24 | genomic DNA from **DH10B ∆*nadA::gfp-kan$^{R}$*** | nadA-gfp-kan for<br>nadA-gfp-kan for | PCR product gel purified, transformed into **DH10B ∆*cpxA::trans* (pKD46)** |
| | 25 | genomic DNAs from previous step | nadA-gfp-kan for<br>nadA-gfp-kan for | diagnostic PCR, product sequenced to confirm correct *nadA* gene disruption and insertion of *gfp-kan$^{R}$* cassette → **DH10B ∆*cpxA::trans* ∆*nadA::gfp-kan$^{R}$* (pKD46)** |
| | | | | **DH10B ∆*cpxA::trans* ∆*nadA::gfp-kan$^{R}$* (pKD46)** cultured at 42 $^{o}$C to remove pKD46 |
| **DH10B *idnK::trans*** | 26 | EB1 genomic DNA | idnK-EB1 for1<br>idnK-EB1 rev1 | **DH10B ∆*rcsC::trans* ∆*nadA::gfp-kan$^{R}$* (pKD46)** cultured at 42 $^{o}$C to remove pKD46 |
| | 27 | PCR products 7+26 | idnK-EB1 for1<br>FRT PGK rev | PCR product gel purified |

| | | | | |
|---|---|---|---|---|
| | 28 | EB1 genomic DNA | idnK-EB1 for2<br>idnK conf rev’ | PCR product gel purified |
| | 29 | PCR products 27+28 | idnK-EB1 final for1<br>idnK conf rev | PCR product gel purified, transformed into **DH10B (pKD46)** |
| | 30 | genomic DNA from #29 | idnK conf for<br>kan rev | diagnostic PCR, product sequenced to confirm correct gene disruption |
| | 31 | genomic DNA from #29 | idnK conf rev<br>kan for | diagnostic PCR, product sequenced to confirm correct gene disruption→ **DH10B** ***∆idnK::trans-kan$^R$*** **(pKD46)** |
| | | | | **DH10B** ***∆idnK::trans-kan$^R$*** **(pKD46)** cultured at 42 ºC to remove pKD46 |
| | | | | **DH10B** ***∆idnK::trans-kan$^R$*** was transformed with pCP20 |
| | | | | **DH10B** ***∆idnK::trans-kan$^R$*** **(pCP20)** was grown at 42 ºC to delete *kan$^R$* and remove pCP20 (*amp$^R$*) → colonies that were sensitive to ampicillin and kanamycin were selected for further characterization |
| | 32 | genomic DNAs from previous step | idnK-EB1 for1<br>idnK conf rev | diagnostic PCR, product sequenced to confirm correct *kan$^R$* gene deletion → **DH10B** ***∆idnK::trans*** |
| | 33 | genomic DNA from **DH10B** ***∆nadA::gfp-kan$^R$*** | nadA-gfp-kan for<br>nadA-gfp-kan for | PCR product gel purified, transformed into **DH10B** ***∆idnK::trans*** **(pKD46)** |
| | 34 | genomic DNAs from previous step | nadA-gfp-kan for<br>nadA-gfp-kan for | diagnostic PCR, product sequenced to confirm correct *nadA* gene disruption and insertion of *gfp-kan$^R$* cassette → **DH10B** ***∆idnK::trans ∆nadA::gfp-kan$^R$*** **(pKD46)** |
| | | | | **DH10B** ***∆idnK::trans ∆nadA::gfp-kan$^R$*** **(pKD46)** cultured at 42 ºC to remove pKD46 |
| **DH10B** ***rcsC ∆356*** | 35 | *spec$^R$* gene cassette | rcsC-specR 356 for<br>rcsC-specR trunc rev | PCR product gel purified, transformed into **DH10B (pKD46)** |
| | 36 | genomic DNA from #26 | rcsC trunc conf for<br>spec rev | diagnostic PCR, product sequenced to confirm correct gene disruption |
| | 37 | genomic DNA from #26 | rcsC trunc conf rev<br>spec for | diagnostic PCR, product sequenced to confirm correct gene disruption→ **DH10B** ***rcsC ∆356-spec$^R$*** **(pKD46)** |

| | | | | |
|---|---|---|---|---|
| | 39 | genomic DNA from **DH10B $\Delta$*nadA::gfp-kan*$^R$** | nadA-gfp-kan for<br>nadA-gfp-kan for | PCR product gel purified, transformed into **DH10B *rcsC* $\Delta$*356-spec*$^R$ (pKD46)** |
| | 40 | genomic DNAs from previous step | nadA-gfp-kan for<br>nadA-gfp-kan for | diagnostic PCR, product sequenced to confirm correct *nadA* gene disruption and insertion of *gfp-kan*$^R$ cassette → **DH10B *rcsC* $\Delta$*356-spec*$^R$ $\Delta$*nadA::gfp-kan*$^R$ (pKD46)** |
| | | | | **DH10B *rcsC* $\Delta$*356-spec*$^R$ $\Delta$*nadA::gfp-kan*$^R$ (pKD46)** cultured at 42 $^o$C to remove pKD46 |
| **DH10B *rcsC* $\Delta$*471*** | 41 | *spec*$^R$ gene cassette | rcsC-specR 471 for<br>rcsC-specR trunc rev | PCR product gel purified, transformed into **DH10B (pKD46)** |
| | 42 | genomic DNA from #41 | rcsC trunc conf for<br>spec rev | diagnostic PCR, product sequenced to confirm correct gene disruption |
| | 43 | genomic DNA from #41 | rcsC trunc conf rev<br>spec for | diagnostic PCR, product sequenced to confirm correct gene disruption→ **DH10B *rcsC* $\Delta$*471-spec*$^R$ (pKD46)** |
| | 44 | genomic DNA from **DH10B $\Delta$*nadA::gfp-kan*$^R$** | nadA-gfp-kan for<br>nadA-gfp-kan for | PCR product gel purified, transformed into **DH10B *rcsC* $\Delta$*471-spec*$^R$ (pKD46)** |
| | 45 | genomic DNAs from previous step | nadA-gfp-kan for<br>nadA-gfp-kan for | diagnostic PCR, product sequenced to confirm correct *nadA* gene disruption and insertion of *gfp-kan*$^R$ cassette → **DH10B *rcsC* $\Delta$*471-spec*$^R$ $\Delta$*nadA::gfp-kan*$^R$ (pKD46)** |
| | | | | **DH10B *rcsC* $\Delta$*471-spec*$^R$ $\Delta$*nadA::gfp-kan*$^R$ (pKD46)** cultured at 42 $^o$C to remove pKD46 |
| **DH10B *rcsC* $\Delta$*705*** | 46 | *spec*$^R$ gene cassette | rcsC-specR 705 for<br>rcsC-specR trunc rev | PCR product gel purified, transformed into **DH10B (pKD46)** |
| | 47 | genomic DNA from #46 | rcsC trunc conf for<br>spec rev | diagnostic PCR, product sequenced to confirm to confirm correct gene disruption |
| | 48 | genomic DNA from #46 | rcsC trunc conf rev<br>spec for | diagnostic PCR, product sequenced to confirm to confirm correct gene disruption→ **DH10B *rcsC* $\Delta$*705-spec*$^R$ (pKD46)** |
| | 49 | genomic DNA from **DH10B $\Delta$*nadA::gfp-kan*$^R$** | nadA-gfp-kan for<br>nadA-gfp-kan for | PCR product gel purified, transformed into **DH10B *rcsC* $\Delta$*705-spec*$^R$ (pKD46)** |

| | | | | |
|---|---|---|---|---|
| | 50 | genomic DNAs from previous step | nadA-gfp-kan for nadA-gfp-kan for | diagnostic PCR, product sequenced to confirm correct *nadA* gene disruption and insertion of *gfp-kan*$^{R}$ cassette → **DH10B *rcsC* Δ*705-spec*$^{R}$ Δ*nadA::gfp-kan*$^{R}$ (pKD46)** |
| | | | | **DH10B *rcsC* Δ*705-spec*$^{R}$ Δ*nadA::gfp-kan*$^{R}$ (pKD46)** cultured at 42 °C to remove pKD46 |
| **DH10B *rcsC* Δ*820*** | 51 | *spec*$^{R}$ gene cassette | rcsC-specR 820 for rcsC-specR trunc rev | PCR product gel purified, transformed into **DH10B (pKD46)** |
| | 52 | genomic DNA from #51 | rcsC trunc conf for spec rev | diagnostic PCR, product sequenced to confirm to confirm correct gene disruption |
| | 53 | genomic DNA from #51 | rcsC trunc conf rev spec for | diagnostic PCR, product sequenced to confirm to confirm correct gene disruption→ **DH10B *rcsC* Δ*820-spec*$^{R}$ (pKD46)** |
| | 54 | genomic DNA from **DH10B Δ*nadA::gfp-kan*$^{R}$** | nadA-gfp-kan for nadA-gfp-kan for | PCR product gel purified, transformed into **DH10B *rcsC* Δ*820-spec*$^{R}$ (pKD46)** |
| | 55 | genomic DNAs from previous step | nadA-gfp-kan for nadA-gfp-kan for | diagnostic PCR, product sequenced to confirm correct *nadA* gene disruption and insertion of *gfp-kan*$^{R}$ cassette → **DH10B *rcsC* Δ*820-spec*$^{R}$ Δ*nadA::gfp-kan*$^{R}$ (pKD46)** |
| | | | | **DH10B *rcsC* Δ*820-spec*$^{R}$ Δ*nadA::gfp-kan*$^{R}$ (pKD46)** cultured at 42 °C to remove pKD46 |
